# Conventional polymers may unintentionally refill in vivo with unassociated drugs

**DOI:** 10.1101/2022.03.21.485180

**Authors:** Kathleen Young, Alan B. Dogan, Christopher Hernandez, Agata A. Exner, Horst A. von Recum

**Author notes:** Author Emails: Kathleen Young, Alan B. Dogan, Christopher Hernandez, Agata A. Exner. Corresponding author address: 10900 Euclid Ave, 220 Wickenden Building, Case Western Reserve University, Cleveland, OH 44106, Horst A. von Recum.

## Abstract

Polymers used as drug delivery devices are ultimately limited by how much drug they can hold; with the device failing if the drug is depleted before the disease is cured. Our lab discovered a means to use thermodynamic driving forces to refill certain classes of polymer after implantation, for additional drug delivery windows. These same, refillable polymers can be used as additives, to provide refilling capacity to classical, non-refillable polymers such as poly(methyl methacrylate) (PMMA). In this paper, we investigated the refilling capacity of another conventional polymer: poly(lactic-co-glycolic acid) or PLGA. We explored both unmodified PLGA implants as well as implants supplemented with polymerized cyclodextrin (pCD) in microparticle form, previously shown to add refillability to poly(methyl methacrylate) (PMMA) implants which were otherwise not refillable. Assessments of in situ forming PLGA implants with and without pCD additives were made, including drug loading capacity in a liquid medium, drug refilling through a tissue-mimicking gel medium, and refilling in *ex vivo* and *in vivo* conditions. Implant cross-sections were imaged via fluorescence microscopy. Drug release from refilled implants, polymer swelling, degradation, phase inversion characteristics were assessed, and drug/monomer computational simulation studies were performed. While generally, the incorporation of cyclodextrin into implants led to significant increases in the amount of refilled drug; unexpectedly, PLGA implants with no incorporated pCD also showed refilling capability. Moreover, in two out of three *in vivo* conditions in rats, PLGA alone showed the potential to refill with comparable, if not greater, amounts of drug than PLGA with pCD incorporated. This contrasts predictions, since PLGA has no specifically designed affinity structure, like pCD does. We theorize that the mechanism for PLGA’s refilling depends on nano-patterning of hydrophilic and hydrophobic molecular domains, giving rise to its affinity-like behavior. The fact that PLGA implants can be refilled with unassociated drugs, gives rise to concerns about the fate of all implants made of poly alpha-hydroxy esters, and likely other polymers as well, and will likely lead to new directions of study such as of unintended polymer / drug interactions.

## Introduction

Traditional drug therapies, such as chemotherapy, still rely on systemic drug delivery throughout the body; this method affects not only the targeted cancer cells but healthy somatic cells, often leading to off-target toxicity. This has led to the development of local drug delivery depots. Localized delivery of chemical therapeutics either through repeated local injections or through solid drug delivering implants can mitigate harmful side effects and are preferred in non-metastatic tumors. However, to maintain therapeutic dosing for extended periods of time, tumor recurrence, or other chronic conditions, either injections must be given frequently or implants need to be replaced.^1–6^

Refillable, reloadable, or replenishable drug delivery devices can help to reduce the frequency of drug administration and of reimplantation procedures by allowing for multiple rounds of drug release from a single implant, offering more attractive solutions for chronic or recurrent conditions. Classic refilling into a drug delivery device typically involves direct injection into an enclosed drug reservoir, and these devices often incorporate design elements for limiting diffusion and controlling drug release, like patterned microchannels and porous membranes.^6–8^ Herein we call this “mechanical refilling”. Newer refillable devices are moving away from reservoir-based designs, instead utilizing chemical affinity interactions between drugs and device materials to refill via local or systemic introduction of drug. Strategies which utilize thermodynamically-favorable affinity for drug delivery include molecular imprinting,^9,10^ bio-orthogonal click chemistry,^11,12^ and cyclodextrin (CD) polymer-based delivery.^2,13^ Herein we call these “thermodynamically refillable devices”.

CD polymer-based delivery takes advantage of molecular affinity between drug molecules and the CD monomer unit. A single CD molecule is a ring of 6-8 glucose units that possesses a hydrophilic exterior and a moderately hydrophobic interior pocket. This structure makes CDs particularly adept at forming inclusion complexes with small hydrophobic molecules, and has led to the widespread use of CDs in pharmaceuticals to solubilize and stabilize labile drugs.^13,14^ When CDs are polymerized into insoluble, durable polymers and utilized as delivery devices, the resulting drug release from such devices is slower, and sustained over a longer period of time compared to non-affinity controls.^2,4,5,10,13,15,16^ Since these inclusion complexes are strong, yet reversible interactions, we have discovered that cyclodextrin polymers can refill with drugs even through complex biological media (e.g. bacterial biofilm, serum biomolecules)^17,18^ and even after initial implantation in vivo.^2^

Our laboratory has previously explored insoluble, polymerized cyclodextrin (pCD) for small hydrophobic drug refilling, and demonstrated its use for many applications including treating cancer, infections, and inflammation.^2,15–20^ Previous work from our lab has also shown that polymers lacking a particular affinity structure (e.g. polymethylmethacrylate (PMMA)) lack drug refilling capabilities.^4^ However, refilling properties could be attained by including an additive polymer with structured affinity (e.g., pCD).^4^ In this paper, we report that while some polymers may not refill (e.g., PMMA), other polymers, which also do not have intentional molecular structure suitable for refilling, may refill nonetheless. Specifically, we observed that the commonly-used polymer, poly(lactic-co-glycolic acid) (PLGA), and perhaps the entire class of poly alpha-hydroxy esters (e.g., PGA, PLLA) can be non-invasively refilled with both local and systemic drug administration. PLGA is one of the most widely used synthetic materials in the clinic. Well-known for its biodegradability and biocompatibility, PLGA has been used to make bioresorbable sutures for nearly 4 decades and is the main component in at least 15 FDA-approved injectable drug delivery products.^21,22^

In this work, we demonstrate the unintended refilling capabilities of PLGA in situ forming implants (ISFIs) and compare them to PLGA ISFIs with 10 wt.% incorporated pCD (a composite formulation that was able to bestow refilling properties to non-refillable PMMA).^4^ PLGA ISFIs are composed of PLGA, a water-miscible organic solvent, a drug, and optional excipients. These implants are administered as injectable fluids and solidify in situ through a process called solvent phase inversion. During phase inversion, the water-miscible solvent in the injected volume exchanges with water in the environment, causing the polymer to precipitate in place as a solid polymer depot.

Depending on the combination of these key components and additives, the phase inversion, drug release, and degradation characteristics of ISFIs can be tuned to the desired profiles.^23–26^ A solvent-inverting ISFIs formulation of PLGA was selected for several practical advantages: ease of manufacture, non-invasive administration, biodegradability, and tunable drug release.^27^ ISFIs has previously not been known to be refillable, and extension of drug delivery from ISFIs could only be performed by injecting more implants or by increasing the implant size.^25^ We applied ISFIs in the context of localized cancer drug delivery to hepatocellular carcinoma, using doxorubicin (DOX) in our published drug model.^28^

In this research, the refilling of this “non-intentionally designed” system (PLGA) was compared to that of an “intentionally designed” system (PLGA-pCD), namely by inclusion of affinity-based pCD polymer microparticles. The intentional design of pCD arises from its monomer structure, which at a fundamental level arises from unique structural and molecular patterning of hydrophobic and hydrophilic regions to achieve significantly increased thermodynamic interactions with small hydrophobic drugs. The reports cited above have demonstrated that carefully and deliberately designed systems can indeed refill with drug non-invasively, or semi-invasively, even after being implanted in the body. However, this paper constitutes a critical report of a polymeric delivery device, which was not intentionally designed for refilling, but was refilled nonetheless with drug *in vitro, ex vivo*, and *in vivo.* This gives rise to the questions of whether PLGA refills with other drugs *in vivo*, and whether poly alpha-hydroxy esters and other implanted polymers in the body also non-intentionally refill with drugs through solvent exchange and favorable thermodynamic interactions.

## Materials and Methods

### Materials

Lightly epichlorohydrin-crosslinked γ-cyclodextrin (γCD) was obtained from Cyclolab, Budapest, Hungary. Poly(DL-lactic-co-glycolic) acid (PLGA) (50:50 2A, Mn 10,000 Da, inherent viscosity of 0.15 dL/g) was obtained from Evonik (Birmingham, AL) and used as received. Doxorubicin hydrochloride (DOX) was obtained from LC Laboratories (Woburn, MA) and used as received. Rat Novikoff hepatoma (N1-S1) cells were obtained from ATCC. N-methyl-2-pyrrolidone (NMP), dimethylsulfoxide (DMSO), ethylene glycol diglycidyl ether (EGDE), 3-(4,5-dimethylthiazol-2-yl)-5-(3-carboxymethoxyphenyl)-2-(4-sulfophenyl)-2H-tetrazolium) (MTS), Iscove’s Modified Dulbecco’s Medium (IMDM), agarose, and phosphate buffered saline (PBS) were obtained from Fisher Scientific (Waltham, MA) and used as received.

### Methods

#### Polymerized Cyclodextrin Microparticle Synthesis

In order to incorporate the refilling capacity of cyclodextrin polymers into the in situ forming PLGA implants, we synthesized pCD microparticles using a previously published protocol.^1^ Briefly, γCD pre-polymer was dissolved in 0.2 M potassium hydroxide to form a 0.25 g/ml solution. EGDE crosslinker was added to the cyclodextrin solution to form a final 1:1.6 (w/v) CD:EGDE solution. The solution was poured into 1% (v/v) solution of Tween 85/Span 85 (24:76 v/v ratio) in light mineral oil at 60 °C and stirred intensely at 700 rpm for 2 h. The resulting particles were washed with increasingly polar solvents to remove residual oil, surfactant, and unreacted crosslinker. Particles were frozen in water and lyophilized for 3 days to obtain a white powder. To prepare drug-loaded microparticles, pCD microparticles were suspended in 10 mg/mL of DOX in DMSO and incubated on an end over end mixer (VWR, Radnor, PA) for 3 days at room temperature. Drug loading was determined both by initial depletion calculation, as well as long-term release studies.

#### Polymer Solution Preparation and Implant Synthesis

In situ forming implants (ISFIs) were synthesized by forming PLGA solutions in water-miscible N-methyl-2-pyrrolidone (NMP), according to previously published protocols.^18^ All polymer solutions were formulated with a 60:40 mass ratio of NMP to combined polymer and drug. Implants containing pCD were made with 10 wt.% pCD at a 60:30:10 mass ratio of NMP:PLGA:pCD. This was the maximum amount of pCD incorporation possible into the ISFIs; above this amount, the solution would not spontaneously form a single solid matrix. Specific mass ratios of implants are listed in **Table 1**. Polymer solution was used within 1 day for implant synthesis. Implants were formed by pipetting 40 uL of polymer solution into 10 mL of PBS in glass vials. The resulting average weight of implants used in this study was 43.3 +/-5.5 mg with average loading of 0.43 mg of DOX (1 wt.%).

**Table 1.**
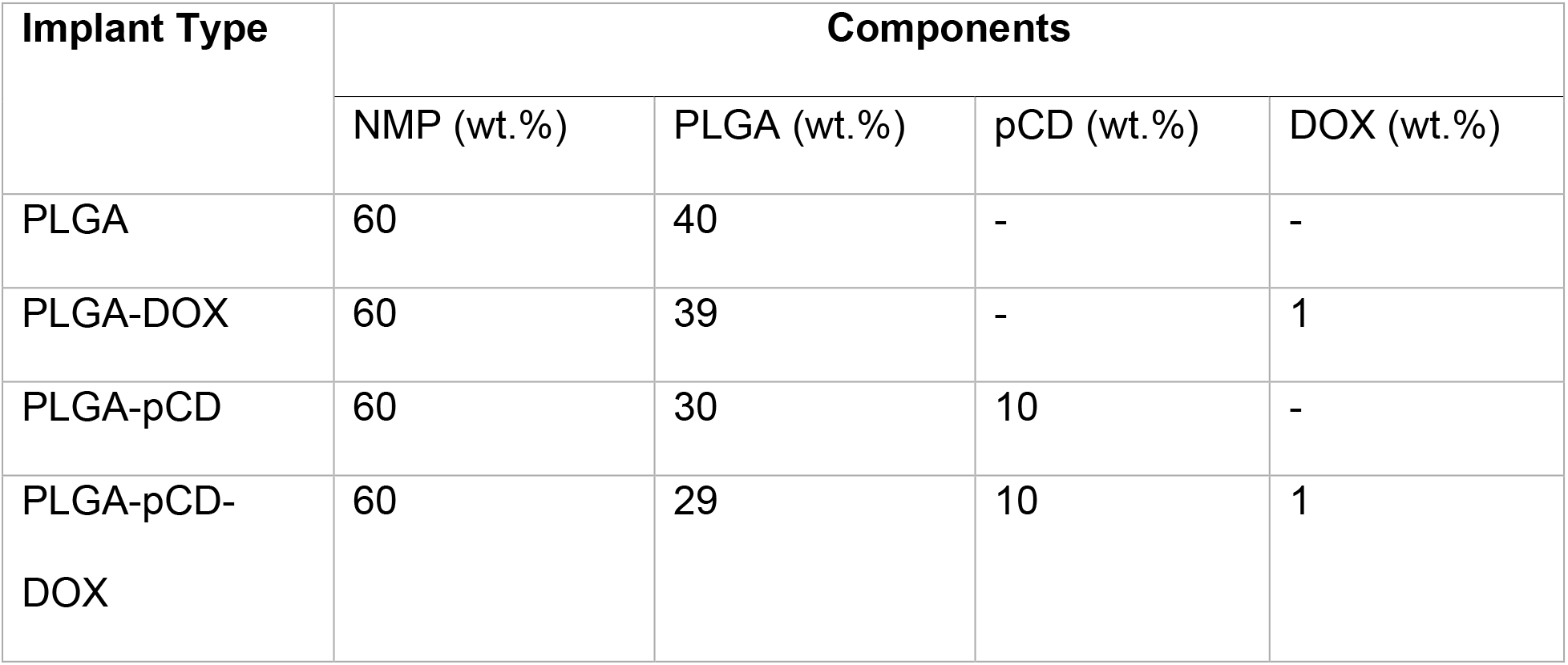
Mass ratios of materials used in polymer formulations. The ratio of solvent to polymer/drug components was kept constant at 60:40. Incorporation of pCD particles was 10 wt.% of the polymer solution, the maximum amount that could be added without exceeding the percolation threshold of the result PLGA-pCD polymer. 1 wt.% DOX is the typical loading amount in DOX-based ISFIs.

**Table 2.** Fold difference in drug loading capacity between PLGA and PLGA-pCD. Difference was calculated as the ratio of DOX loaded into PLGA-pCD to DOX loaded into PLGA.

| Drug Loading Solution Concentration, DOX/PBS (mg/mL) | PLGA DOX loaded (ug/mg) | Standard Deviation | PLGA-pCD DOX loaded (ug/mg) | Standard Deviation | Fold Difference |
| --- | --- | --- | --- | --- | --- |
| 0.2 | 0.639 | 0.211 | 0.896 | 0.104 | **1.402** |
| 1 | 1.745 | 0.323 | 2.443 | 0.533 | **1.400** |
| 5 | 2.016 | 0.767 | 5.594 | 0.533 | **2.775** |

#### Drug Loading Capacity *In Vitro*

To evaluate the maximum amount of drug that could be loaded into “empty” implants, implants without drug were placed in high concentration drug loading solutions, and the amount of loaded drug was determined by PLGA dissolution and solvent extraction. Briefly, implants without drug (PLGA-only, PLGA-pCD) were synthesized and stored on an incubated shaker at 37 °C and 100 rpm for 3 days prior to use. Drug loading solutions were made at concentrations of 0.2, 1, and 5 mg/mL of DOX in PBS. Implants were incubated in loading solutions at 37 °C and 100 rpm for 3 days, removed from the solutions to end loading, rinsed briefly in DI water, then patted dry with tissue (Fig. 1A). To quantify the amount of loaded DOX, implants were dissolved in DMSO, and overall drug fluorescence was quantified on a multimode microplate reader (Synergy, H1 Microplate Reader). Fluorescence was compared to a standard curve of known DOX concentrations to convert fluorescence to drug mass.

**Figure 1.**
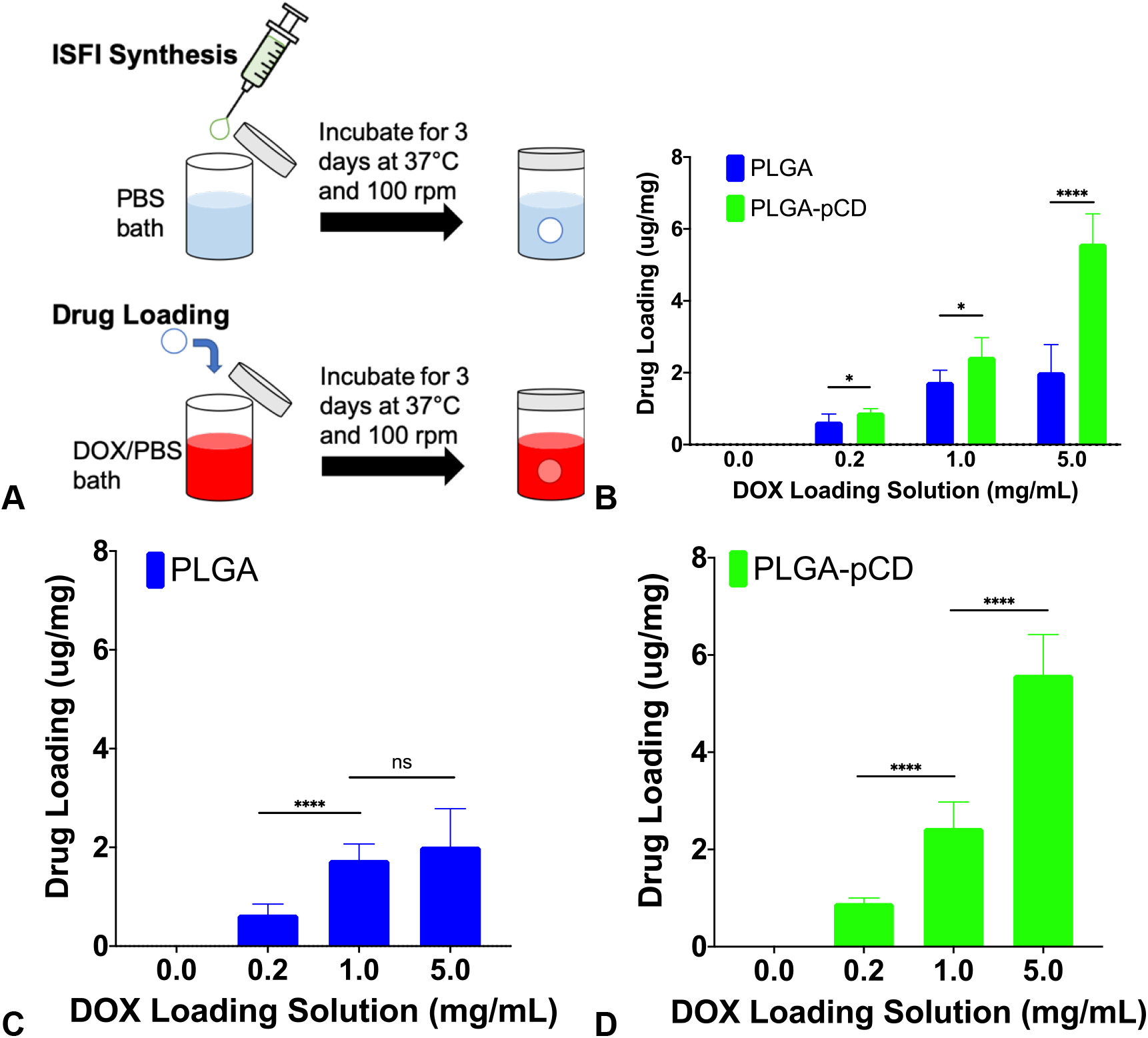
Quantification and comparison of DOX loaded into implants by immersing in concentrated drug solution. **A**) Schematic depiction of ISFIs synthesis and drug loading procedures. To determine maximum loading, drug-loaded implants were dissolved in DMSO, and drug content was analyzed using fluorescence spectroscopy. Drug loading capacity is reported as loaded drug (ug) divided by initial polymer weight (mg). **B**) Comparison of drug loading capacity of both PLGA and PLGA-pCD. **C**) Drug loading capacity of PLGA-only. PLGA drug loading capacity essentially saturated near 2 ug/mg with a drug loading solution of 1 mg/mL DOX in PBS; no significant difference between drug loading was observed at 1 and 5 mg/mL. **D**) Drug loading capacity of PLGA-pCD. Drug loading into PLGA-pCD continued to increase significantly beyond 1 mg/mL, and it is likely that the drug loading capacity of PLGA-pCD will saturate when an even higher concentration of drug loading is used. Two-sample Student’s t-test was used for statistical analysis; n=3 for control (0 mg/mL); n=6 for experimental groups. * for p-value<0.05, **** for p-value<0.0001. Error bars depicted are standard deviation.

#### Drug Refilling *In Vitro* Through Tissue-Mimicking Agarose Medium

To evaluate the capacity of a drug-depleted device to refill in the body through tissue, we utilized a previously published tissue-mimicking agarose phantom assay.^17,29^ Implants without drug (PLGA-only and PLGA-pCD) were synthesized and stored in PBS on an incubated shaker for 3 days prior to use. Agarose gels were made using 0.075% (wt/vol) agarose/PBS solution. Agarose solution was added to 6 well plates and allowed to cool. Implants were placed on the agarose, and agarose solution was added to encapsulate the implants. A biopsy punch was used to form a 6 mm diameter well in the center of each agarose gel. Drug solution (1 mg/mL DOX in PBS) was added to the wells. The plate was sealed with Parafilm and placed on an incubated shaker at 37 °C and 100 rpm for 3 days. The progression of DOX refilling throughout the gels was monitored using the area scan function of the same multimode microplate reader as described above. After refilling was completed, implants were excised from the gels. The relative amounts of refilled DOX were determined by fluorescence macro-imaging (Maestro In Vivo Fluorescence Imaging System, Cambridge Research & Instrumentation, Inc). To assess the depth of refilling, cross-sections of refilled implants were viewed in two ways. First, implants were frozen in liquid nitrogen and then freeze-fractured to expose domain of refilled, colored drug. Photos were taken of the fractured cross-sections. In a second method, implants were frozen in OCT medium and sectioned on a HM 505 E microtome cryostat (Microm International GmbH, Walldorf, Germany) and imaged via the same fluorescence macro-imaging technique as stated above.

#### Drug Refilling Through Liver Tissue, *Ex Vivo*

For this study we also developed an ex vivo refilling assay that can be used as a model to assess refilling through tissues and extracellular fluid. A modified agarose phantom assay was used similar to above, except that a large piece of tissue (e.g. liver) was embedded in the drug diffusion region. Fresh pig liver was obtained from animals otherwise sacrificed during unrelated surgical training at the Case Western Reserve School of Medicine Animal Resource Center. The liver was cut into approximately 3 by 3 by 1 cm rectangular pieces and stored in at 4 °C for 2 days to exsanguinate. Implants without drug were synthesized and stored in PBS on an incubated shaker for 3 days prior to use. Implants were removed from PBS and blotted briefly on tissue to remove excess liquid on implant surfaces prior to use. Implants were embedded into liver pieces by cutting pockets into the tissue with surgical scissors and placing pre-formed implants into the pockets. Then the liver / ISFIs complex was fully embedded in agarose in the wells of 6-well plates. A competitive assay was used, in which 2 implants were embedded into a single piece of liver: both an experimental (PLGA-pCD) and a control (PLGA-only) implant (**Fig. 4A**). The implants were positioned approximately 1.5 cm from each other. Drug solution (100 uL of 1 mg/mL DOX in PBS) was injected into the liver equidistant from each of the implants. The plates were sealed with Parafilm and placed on an incubated shaker at 37 °C and 100 rpm for 3 days. After refilling was completed, implants were excised from the gels and liver, rinsed briefly in PBS to remove excess unbound drug, and blotted dry on tissue. The amount of refilled drug was quantified by dissolving implants in DMSO and then measuring the amount of DOX via fluorescence spectroscopy, as described above.

#### Drug Release from Refilled Implants, *In Vitro*

To understand the refilled implants’ subsequent release profile, we carried out release studies from PLGA and PLGA-pCD implants refilled via diffusion of DOX through agarose tissue phantoms. Implants were synthesized and refilled according to the tissue-mimicking agarose phantom assay described in Drug Refilling in vitro. Refilled implants were removed from agarose gels and placed in 10 mL PBS on an incubated shaker at 37 °C and 100 rpm. For sampling, 1 mL of release solution was sampled and replaced with 1 mL fresh pre-warmed PBS. Mass of released DOX was determined by measuring the fluorescence (excitation: 498 nm, emission: 590 nm) in the solution samples and referenced to a standard curve of known masses on a Synergy H1 Microplate Reader (BioTek, Winooski, VT).

#### Drug Refilling, *In Vivo*

In order to assess refilling capacity in vivo, a proof-of-concept refilling study was carried out in rats. All animal experiments were performed in accordance with relevant guidelines and regulations of Case Western Reserve University Institutional Animal Care and Use Committee. Wild-type male Sprague-Dawley rats were injected with 50 uL of ISFIs solution through a 21-gauge needle. Mass of the ISFIs syringe was measured before and after injection to ensure equivalent ISFIs injections. A control (PLGA) and an experimental (PLGA-pCD) ISFIs was implanted in each rat. Implants were created by injecting polymer solution subcutaneously, intramuscularly, and into the liver, to assess refilling in different tissue environments (n=2 per injection location). Three days after implant injection, a refilling dose of Doxil was administered subcutaneously near each subcutaneous implant, intramuscularly near each intramuscular implant, and systemically by tail vein injection for liver implants. To end refilling after administration of the refilling dose, animals were euthanized following procedures described in the approved animal protocol 3 days after refilling dose. Implants were harvested, and then cut in half longitudinally for imaging. Levels of refilled DOX were determined via fluorescence macro-imaging as previously described.

#### Molecular Docking Simulations Between Drug and Monomers

Molecular docking simulations were performed to assess theoretical binding affinities between the drug DOX and the components of PLGA and PLGA-pCD ISFIs: PLGA monomers, lactic acid (LA) and glycolic acid (GA); and gamma-CD monomers, using the virtual screening software AutoDock Vina in Pyrx.^30^ An sdf file of the guest molecule (DOX) and pdb files of the host molecules (LA, GA, gamma-CD) were uploaded to Pyrx. The guest molecule was minimized and then converted to an AutoDock Ligand pdbqt file. The host molecules each were converted to macromolecule files. The Vina Wizard was used to perform the docking simulation. Briefly, a pair of guest and host molecules were selected, the macromolecule space was maximized, and all other settings were left on their defaults. The relative binding energies of different host/guest combinations were then compared to evaluate binding affinity strength.

#### Statistical Analysis

Data are represented as mean with standard deviation. Statistical significance was defined as p < 0.05. Two-sample Student’s t-test was used for statistical testing; further specifications and alternative tests used are noted in the results and figure captions.

## Results

### Drug Loading Capacity *In Vitro*

To assess the drug loading capacity of PLGA and PLGA-pCD ISFIs, which would give us an idea of the greatest potential of these polymers to refill with drug, we performed an *in vitro* drug loading experiment, exposing drug-free implants to concentrated drug solutions (0.2 mg/mL, 1 mg/mL, 5 mg/mL, at n=6) to facilitate drug loading (**Fig. 1A**). Implants stored in drug-free PBS were used as the negative control (n=3). The presence of pCD significantly increased drug loading capacity, with significantly more drug loaded into PLGA-pCD across all loading conditions (**Fig.1B**). Student’s two-sample t-test was use for hypothesis testing. At loading concentrations of 0.2 and 1 mg/mL, PLGA-pCD loaded with significantly greater amounts of DOX (p=0.023 and p=0.021 respectively). The greatest difference between drug loading into PLGA and PLGA-pCD was seen at the highest DOX concentration of 5 mg/mL, where PLGA-pCD loaded with approximately 2.8 times more DOX than PLGA (p=0.000015). No significant difference was found between PLGA and PLGA-pCD implants at 0 mg/mL, the control refilling concentration. Drug loading into PLGA-pCD continued to increase significantly with each increase in DOX concentration (**Fig. 1D**). However, loading into PLGA did not increase significantly beyond 1 mg/mL (p=0.44, non-significant) (**Fig. 1C**), indicating that drug loading at higher concentrations was close to becoming saturated. Overall, incorporation of pCD enhanced the drug loading capacity of PLGA-based ISFIs.

### Drug Refilling Through Simple, Tissue-Mimicking Agarose Model

We performed in vitro drug refilling experiments to assess the refilling capacity of ISFIs when refilled via local administration of DOX through a tissue-mimicking medium. Implants were synthesized, precipitated for 3 days, and then were embedded in agarose gels which acted as a tissue phantom (**Fig. 2A**). The agarose was inoculated with DOX in PBS, which then diffused throughout the gel and refilled into the embedded polymers. After 3 days, implants were harvested and evaluated for the total amount of refilled drug as well as the depth of refilling. The relative amounts of refilled drug in intact implants were quantified via fluorescence macro-imaging (**Fig. 2B**), showing that PLGA-pCD refilled approximately 1.8 times more drug compared to PLGA (p=0.002, Student’s two-sample t-test). To evaluate depth of refilling, implant cross-sections were exposed via freeze fracture (**Fig. 3A**). Photos of freeze-fractured implants indicate that refilling of PLGA was limited to the surface, and that DOX was able to refill further into the interior of PLGA-pCD. Refilling depth was also assessed by fluorescence imaging of implant cryosections, which also showed greater depth of refilling in PLGA-pCD (**Fig. 3B**).

**Figure 2.**
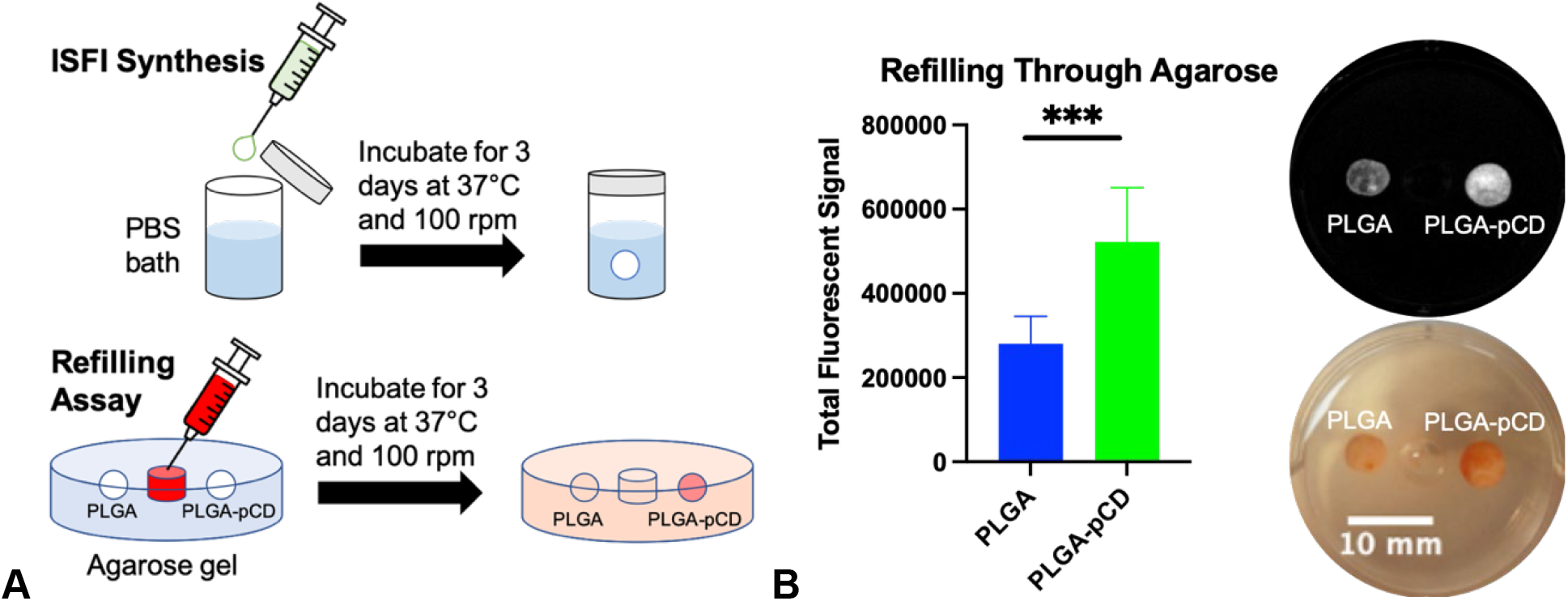
Drug refilling was performed in tissue-mimicking agarose phantoms for 3 days. **A**) Schematic of the tissue-mimicking agarose assays. Implants were synthesized 3 days prior to embedding into agarose, and refilling proceeded over 3 days. The agarose gels depicted represent a single well of a 6-well plate. **B**) Overall fluorescent signal was approximately 1.8 times greater in PLGA-pCD. Insets: representative fluorescent and photo image of a single agarose gel post-refilling. Two-sample Student’s t-test was used for statistical analysis. *** indicates p value < 0.005.

**Figure 3.**
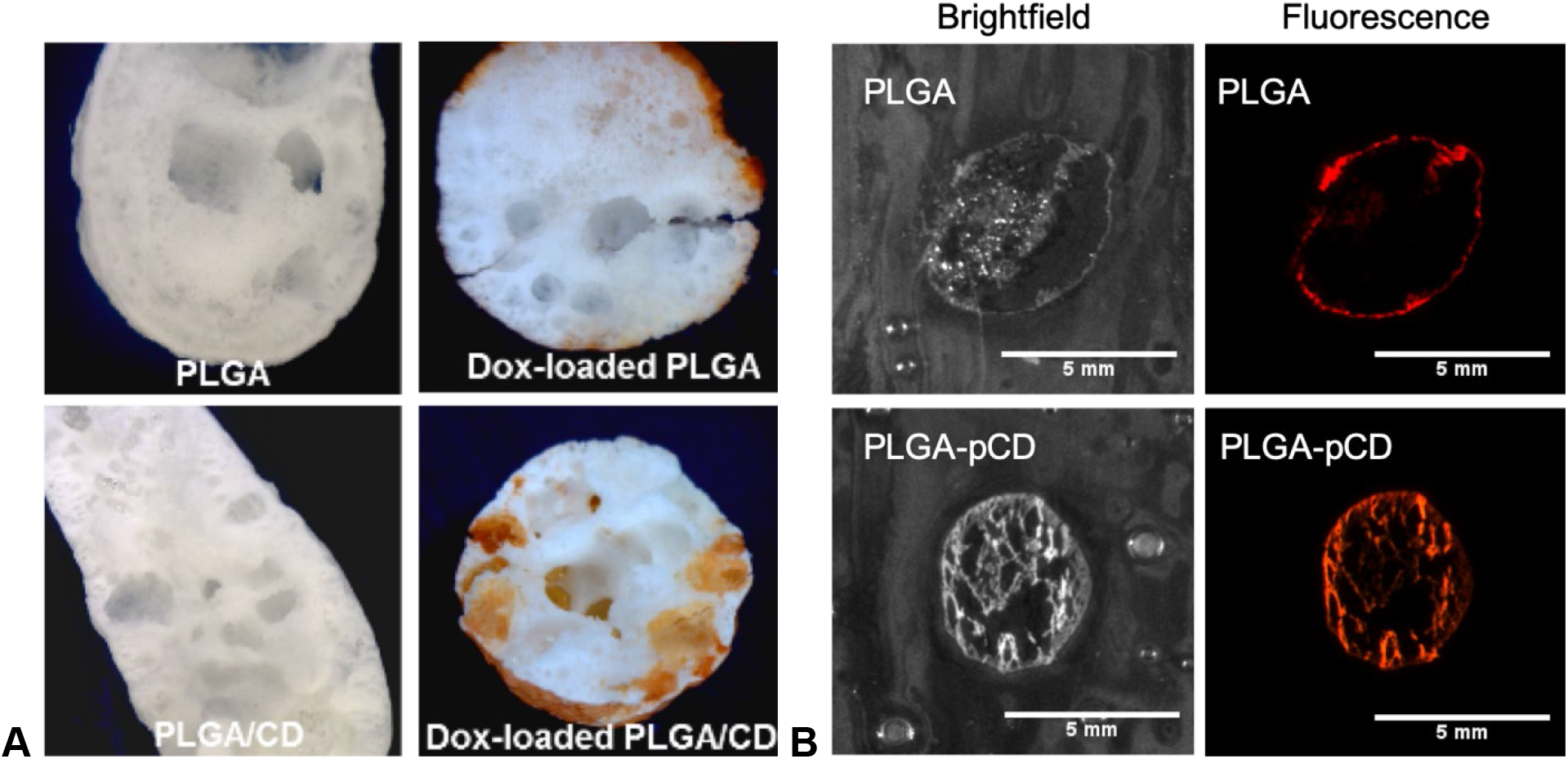
Differences in drug refilling depth between PLGA and PLGA-pCD. **A**) PLGA and PLGA-pCD implants were freeze-fractured to view cross-sections. Domains of refilled drug were visible in the interior of PLGA-pCD implants as dark red-orange spots, whereas drug refilled primarily on the surface of PLGA. **B**) Brightfield and fluorescent images of representative cryosections of refilled implants showing differences in depth of refilling. Similar to the freeze fractured cross-sections, much greater depth of refilling was observed in PLGA-pCD compared to PLGA.

### Drug Refilling through liver tissue *ex vivo*

Due to the heterogeneity of hepatic tissue, we aimed to see if an inhomogeneous, challenging diffusion medium would impact polymer refilling. We carried out *ex vivo* drug refilling assays through bulk liver tissue (which was embedded within agarose) to assess drug refilling through a more complex, biological medium (**Fig. 4A**); namely, of diffusion through tissue and extracellular fluid. We found that both PLGA and PLGA-pCD depots refilled to some capacity: 2.1 and 3.7 times greater than the corresponding negative controls, respectively. However, the only statistically significant difference was between refilled and control (not refilled) PLGA pCD (p=0.0021, Student’s two-sample t-test). Refilled PLGA-pCD refilled 1.8 times more than refilled PLGA, however this difference was not statistically significant (p=0.2452). Negative controls contained notable baseline fluorescence which might have been attributed to autofluorescence of the polymer ISFIs and components of the liver tissue which could not get washed out. PLGA-pCD showed significantly higher autofluorescence compared to PLGA (p=0.0222) (**Fig. 4B**).

**Figure 4.**
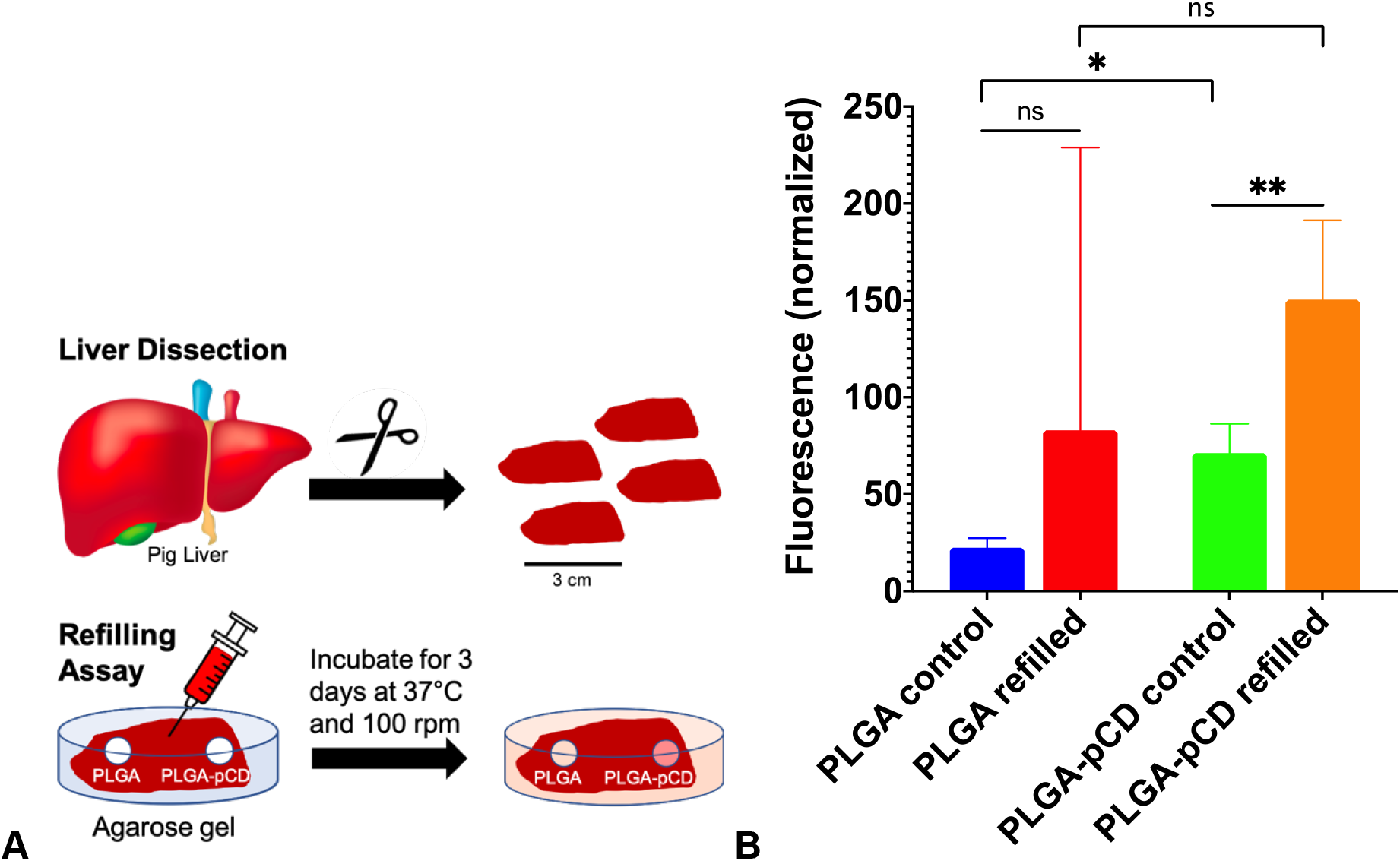
Refilling in an *ex vivo* model utilizing pig liver tissue was used to compare refilling ability. **A)** Schematic of how the *ex vivo* liver refilling assay was carried out (see methods). **B**) Relative fluorescence of PLGA and PLGA-pCD ISFIs, normalized by implant weight, after refilling. Error bars given as standard deviation, n=3. ns=nonsignificant. * indicates p-value<0.05. ** indicates p-value<0.01.

### Drug Refilling *In Vivo*

In order to assess the potential of PLGA-pCD ISFIs to refill with drug *in vivo*, a proof-of-concept refilling study was carried out on ISFIs which were implanted into various locations in rats. Both PLGA and PLGA-pCD were used in the same animal, and implants were refilled by local or systemic administration of DOX. **Fig. 5A** shows the fluorescence intensities of refilled implants recovered from animals after the refilling procedure. Subcutaneously injected PLGA-pCD combined with local subcutaneous refilling exhibited roughly 4 times the amount of average fluorescence compared to PLGA (**Fig. 5B**). In contrast, for intramuscular ISFIs combined with local refilling, PLGA seemed to exhibit roughly 1.3 times more fluorescence than PLGA-pCD (**Fig. 5C**). In a third model, ISFIs were injected into livers. Refilling occurred in these animals through a drug dose administered systemically via tail vein. In this scenario, PLGA also seemed to show more refilling than PLGA-pCD (**Fig. 5D**); however, these differences were not statistically significant, due to the small sample size.

**Figure 5.**
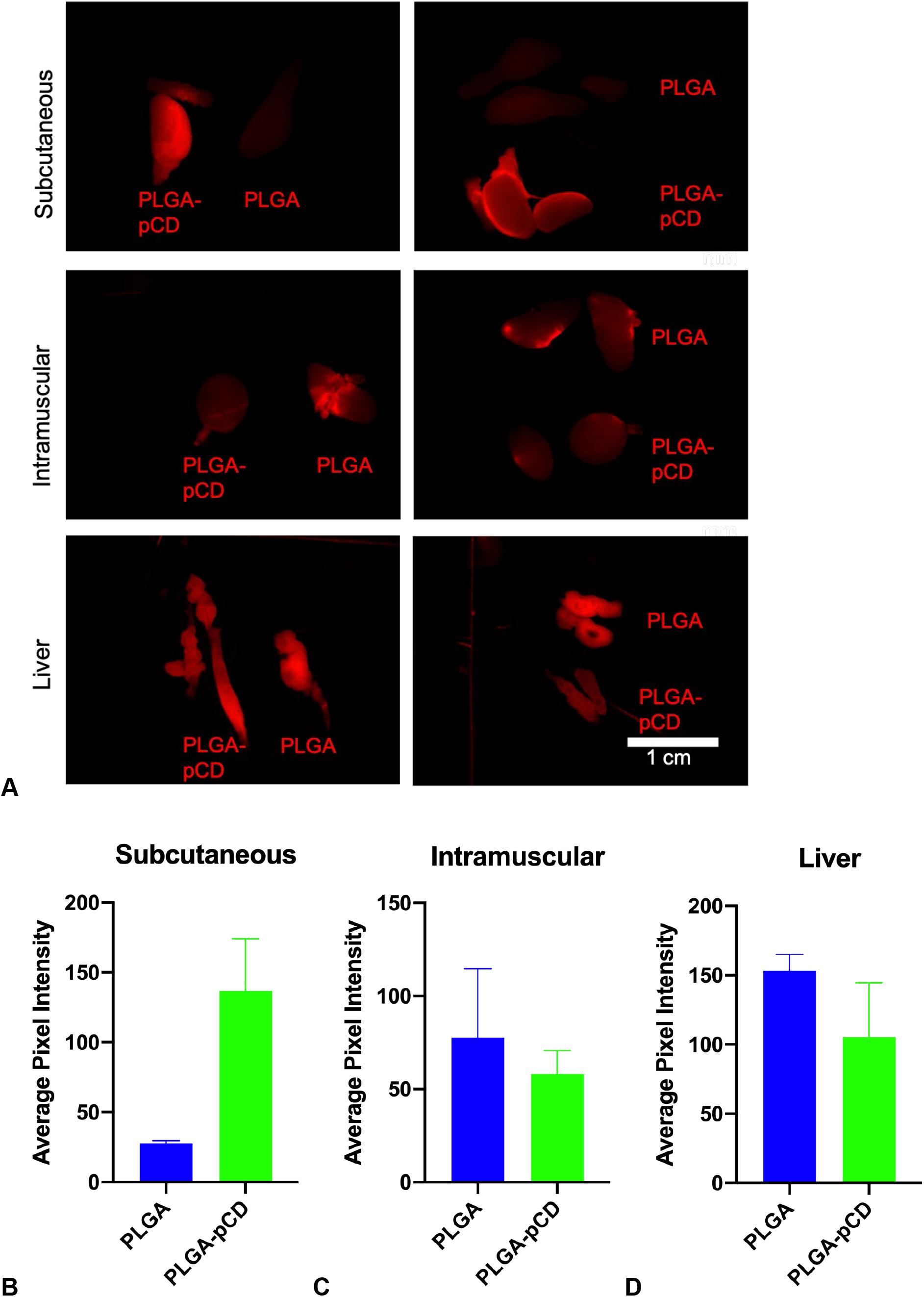
Fluorescence intensity of DOX in implants refilled *in vivo*, n=2. Implant halves are placed next to each other. **A**) Subcutaneous implant injection and local injection of refilling DOX. Intramuscular implant injection and local injection of refilling DOX. Implant injection in liver vasculature and systemic injection of refilling DOX. Relative amounts refilled given as average fluorescent pixel intensity of **B**) subcutaneous (refilled locally), **C**) intramuscular (refilled locally), and **D**) liver implants (refilled via tail vein injection).

### Drug Release from Refilled Implants

To understand the subsequent release profile from refilled implants, we carried out release studies from PLGA and PLGA-pCD refilled with DOX in agarose tissue phantoms. The release profiles from both types of polymers were similar (no significant difference) (**Fig. 6**).

**Figure 6.**
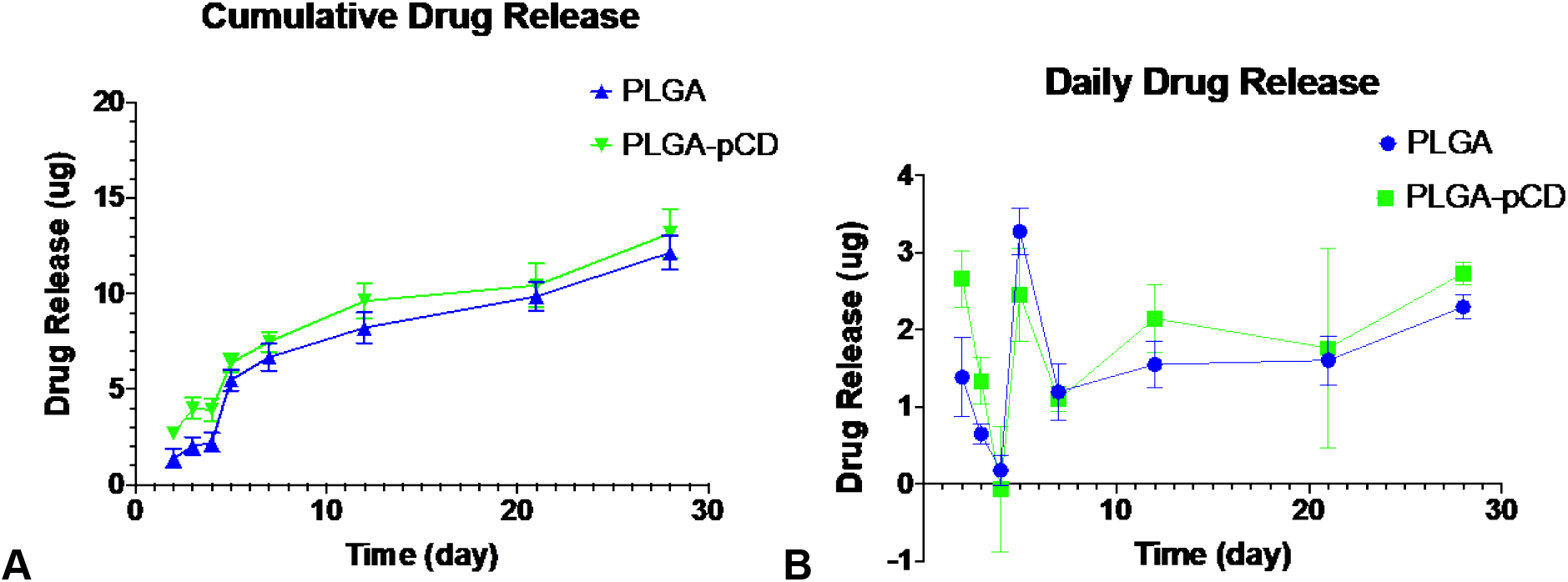
Implants were first refilled with DOX through an agarose tissue phantom. Subsequent DOX release was measured. **A**) Cumulative DOX release over 28 days. **B**) DOX release depicted as the daily amount released. Similar release profiles were observed from both refilled PLGA and PLGA-pCD (non-significant difference).

### Binding Affinity from Molecular Docking Simulations

The unpredicted affinity between PLGA and DOX (Fig. 5) led us to perform docking simulations to assess theoretical affinity between PLGA monomers, LA and GA, and the drug DOX in the program PyRx. This was to see whether individual components of PLGA contributed to its affinity with DOX, since drug affinity with pCD is due to the interaction between DOX and the individual CD monomers of pCD. Simulations showed strong binding affinity between gamma-CD monomer and DOX at −7.4 kcal/mol.^2^ However, simulations showed no particular affinity between PLGA monomers and DOX (**Table 3**). GA and LA have nonzero affinity with CD, but the magnitude of these affinities are almost an order of magnitude lower than that between DOX and gamma-CD. Polymers with affinity constants of these magnitudes in our hands have showed negligible refilling.

**Table 3.** Simulated binding affinity between DOX and monomers of PLGA (glycolic and lactic acid), and pCD. DOX was used as the guest molecule. LA, GA, and gamma-CD were used as host molecules. Binding affinity values given on a log scale.

| Host | Guest | Binding Affinity<br>(kcal/mol) |
| --- | --- | --- |
| Glycolic Acid | DOX | -1.9 |
| Lactic Acid | DOX | -2.3 |
| γCD | DOX | -7.4 |

## Discussion

We investigated the impact of pCD incorporation on the drug loading and refilling behavior of PLGA-based ISFIs, examining drug refilling *in vitro, ex vivo*, and *in vivo*, as well as drug release from refilled materials. Other ISFIs properties were also assessed: swelling, degradation, and initial drug delivery (Supplementary Materials). During this investigation, we found that PLGA-pCD refilled with more drug as we expected, due to the well-demonstrated capacity of DOX to form inclusion complexes with CD.^2^ Specifically, under in vitro (**Fig. 1-3**) and ex vivo (**Fig. 4B**) contexts, the incorporation of pCD microparticles significantly increased drug loading and refilling capacities of PLGA-based ISFIs. In particular, PLGA-pCD refilled robustly through dense, complex tissue matrix. PLGA, however, was also observed to refill with substantial amounts of drug including under some *in vivo* conditions. In a proof-of-concept study, PLGA exhibited refilling capacity *in vivo* comparable to PLGA-pCD (**Fig. 5C-5D**). Of note, this distinction was best found in PLGA ISFIs implanted into the liver.

Drug refilling efficiently into pCD compared to other non-affinity polymers has been demonstrated for several combinations of pCD and small hydrophobic drugs.^2^ A past study evaluating refilling in PMMA with and without CD showed that PMMA by itself showed essentially no refilling.^31^ The study contained herein represents the first time refilling has been demonstrated for a polymer that has no specifically-designed mechanism for refilling (PLGA). This unanticipated finding could be transformative, since there are many medical uses of PLGA, poly alpha-hydroxy esters (e.g. PGA and PLLA) and potentially many other polymers used clinically which could be impacted by accidentally refilling with drugs unrelated to the implant function.

While affinity between drug and pCD monomer units (CDs) drives the affinity-based release and refilling in the intentionally-designed systems,^2,5^ we know from computational modeling that the affinities between PLGA monomers (LA and GA) and the drug tested here (DOX) are negligible (**Table 3**). We theorize that it is the amphiphilic nature of PLGA that makes refilling possible. Namely, nano-domains of aggregated hydrophobic monomer units (LA) will exist in a sea of hydrophilic monomer units (GA), with these amphiphilic domains somewhat recapitulating the amphiphilic structure of cyclodextrin (**Fig. 7**), and will passively facilitate the sequestration of hydrophobic drug molecules into the hydrophobic nanodomains. However, while this theory can explain how PLGA is able to refill with drug at all, it does not explain why PLGA showed comparable, if not better, levels of refilling *in vivo*.

**Figure 7.**
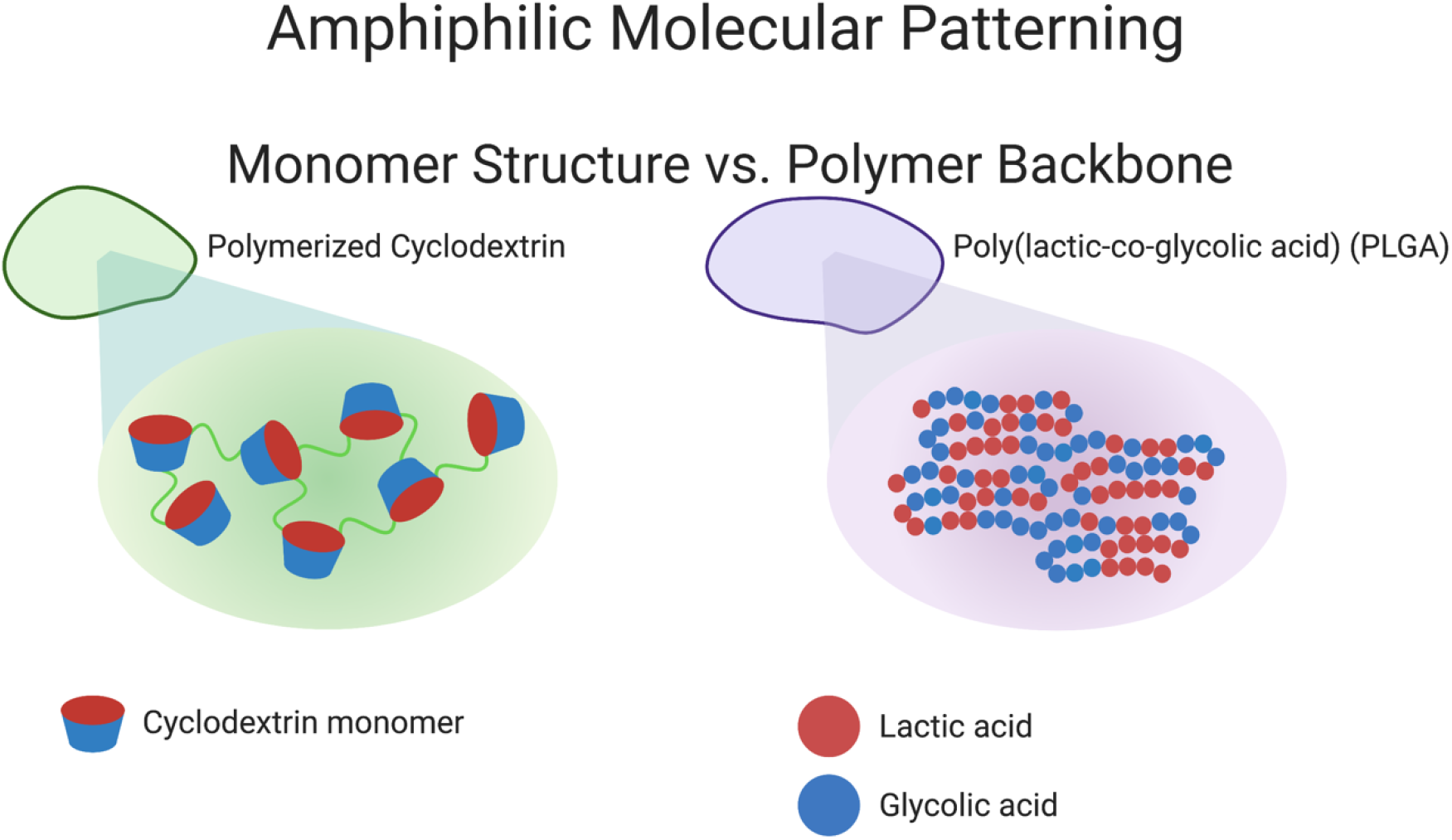
Schematic of “Amphiphilic Molecular Patterning” in pCD vs. PLGA, showing two different ways in which amphiphilic patterning can arise. In pCD on the left, CD monomers produce the patterning effect, with hydrophobic CD pockets surrounded by their hydrophilic exteriors. The interactions between drug molecules and individual CD monomers dictate the level of affinity seen in the polymer. In the random copolymer of PLGA on the right, patterning is likely more irregular due to the random incorporation of hydrophobic LA monomers in the polymer backbone. The random configuration of larger groupings of LA appear as the polymer chains pack, producing relatively hydrophobic regions surrounded by more hydrophilic GA monomers. Figure created with BioRender.com.

We hypothesize that two factors related to the environment could be influencing PLGA refilling behavior *in vivo*. For one, *in vivo* injection locations are known to affect ISFIs physical structure features, like pore size and implant shape,^32^ which can significantly influence drug refilling. ISFIs used for *in vitro* and *ex vivo* experiments were pre-formed in PBS in order to achieve relatively uniform spherical shapes and pore size distribution. Conversely, for *in vivo* experiments, ISFIs were injected and therefore produced heterogeneous shapes and pore sizes dependent on injection location. Second, the comparable refilling of PLGA may have occurred because pCD refilling to its full capacity is dependent on the duration of exposure to a drug source. PLGA-pCD refilled more efficiently under conditions in which the refilling drug dose was effectively contained near the implant and underwent minimal clearance/elimination from the system, i.e. *in vitro, ex vivo*, and subcutaneously *in vivo* (**Fig. 1-4, 5A**). *In vitro* and *ex vivo* refilling assays proceeded over 3 days to allow drug diffusion to reach equilibrium, with no drug depletion by the environment, giving an idea of the maximum amount of drug refilling given the conditions used. *In vivo*, the retention and clearance of small molecule drugs like DOX in a particular tissue or organ depends on cardiac output and blood perfusion rate. The subcutaneous/skin space has the lowest cardiac output and blood perfusion rate of the 3 locations used^33^, and thus would have the most drug retention over the duration of the refilling period. This is supported by our data, in which refilling of PLGA-pCD *in vivo* was higher subcutaneously and lower in the muscles and liver. Implants injected intramuscularly and into the liver likely had a briefer period of exposure to DOX from clearance due to higher blood flow and metabolic clearance. Within a short timescale, PLGA and PLGA-pCD ISFIs showed similar amounts of DOX refilling.

In regard to the release profile from refilled implants, there was no significant difference between PLGA and PLGA-pCD. This indicates that for the period of the drug release study, drug solubility governed release. Given that PLGA-pCD refills with a greater amount of DOX, we hypothesize that PLGA-pCD will continue to release DOX even after PLGA has fully degraded. We also quantified drug release, first from PLGA-pCD implants in which DOX was loaded only into pCD, and then the drug-loaded pCD were incorporated into PLGA polymer solution (**Fig. S3**). In these implants, DOX was not initially distributed throughout the PLGA phase, which significantly decreased burst release and slowed down subsequent drug release. For ISFIs which had DOX incorporated directly into the polymer solution, rather than only into pCD microparticles, PLGA-pCD ISFIs experienced greater burse release (due to accelerated phase inversion) and greater cumulative drug release compared to PLGA ISFIs (**Fig. S4**). An ideal formulation of DOX-delivering ISFIs should have DOX pre-loaded into pCD microparticles as well as directly incorporated into polymer solution. This would maximize both drug release duration as well as cumulative drug released, in addition to providing an immediate bolus of DOX to suppress existing cancer cells in the vicinity.

We acknowledge several limitations of this work. Regarding the type of drug delivery implant selected, drug release from diffusion-governed drug delivery devices have limited drug penetration depth through tissue, limiting therapeutic effects to within a few millimeters of the device.^34^ However, it is known that PLGA implants with increased swelling (generally more swelling with lower molecular weight polymers) experience greater pressure from the surrounding tissue environment, creating compressive forces that induces convective transport of solvent and drug from implants. This may even increase tissue penetration depth.^32^ PLGA-pCD is likely to exhibit this effect to a significantly greater degree than PLGA alone, as it showed significantly more swelling (**Fig. S1**). This is due to the hydrophilicity and hydrogel-like nature of pCD increasing water uptake and retention. The results of this study are not universal for all ISFIs; a single PLGA type and CD type were used in this study to simplify analysis. Gamma-CD was selected due to its demonstrated higher affinity for DOX than other cyclodextrins.^2^ For a different disease, another combination of drug, PLGA, and CD may need to be selected to obtain a more appropriate drug release profile. Finally, only a single round of refilling was used to assess drug loading and reloading efficiency. Ideally, a refillable device would be refilled more than once, and its refilling efficiency for multiple consecutive refills would need to be assessed to determine whether that efficiency changes over time.

To address these limitations, future work should include investigation of multiple rounds of drug refilling, as well as the use of other combinations of PLGA and pCD, in rigorous animal studies. The propensity for PLGA to refill could be more advantageous with slower degrading formulations with higher molecular weight, and/or higher ratio of LA to GA. Strategies involving refilling into pCD-based devices should include methods of increasing drug retention at local injection sites to improve refilling. Refillable ISFIs can potentially be used for cancer immunotherapy, a field in which recent work suggests that localized therapies may be more effective than systemically administered ones in certain cases.^35^ This study also suggests that molecular docking simulations may be insufficient to capture affinity interactions between drugs and PLGA, and possibly other polymers. To better predict interactions between drugs and polymers, there is a need to better similar actual interactions between drugs and polymers, which will necessitate further development of molecular modeling techniques.^36^

## Conclusions

It had been assumed that most polymers do not refill with drug after implantation and thus are single-use devices with finite therapeutic lifetime. Previous work in our lab demonstrated that due to the capacity of monomer/drug complex formation, pCD is capable of affinity-based refilling in the presence of exogenous drug, refilling a great deal more than control, non-affinity polymers (dextran, PMMA). These works also showed that previously non-refillable polymers could be made refillable through the addition of the affinity polymer pCD. ^2,10,17^ This paper shows that PLGA, one of the most commonly-used biocompatible and biodegradable polymers, is also capable of refilling with drug and in certain condition at levels comparable to that of polymers with pCD additives, even though PLGA has no specific molecular design to be refillable. The incorporation of pCD into PLGA increases drug loading and drug refilling capacity compared to PLGA alone, yet proof-of-concept *in vivo* studies showed that PLGA could potentially have even higher levels of refilling than PLGA-pCD in certain conditions.

Based on this work, the interactions of the biomedical polymer PLGA as well as other poly alpha-hydroxy esters with commonly-used drugs should be reassessed. Additionally, other commonly used polymers should be investigated for their unintended refilling with unassociated drugs. Many patients receiving PLGA-based implanted biomaterials (e.g. elderly) are likely to also be taking a cocktail of drugs for other conditions, and as such there might be unwanted affinity between their PLGA device and these other drugs, even in the presence of biological molecules.^18^ This refilling could result in a range of unwanted effects, including sequestering of drug away from where it is needed, as well as re-delivery of drug where it is not needed (e.g. a chemotherapy drug in a vascular stent). Lastly with degradable polymers such as PLGA, refilling of unanticipated drug could impact the degradation rate of the polymer.

While the FDA is a well-respected agency for monitoring human use of drugs (CDER) and implants (CBER), its bicameral approach may not be optimal for evaluating and predicting unassociated drug-implant interactions. Future work should focus on the high-throughput assessment of the ability of commonly implanted biomaterials to non-specifically absorb a broad range of small hydrophobic pharmaceuticals.

## Supporting information

Supplemental Methods and Results

## Acknowledgements

Research reported in this publication was supported by the National Science Foundation Graduate Graduate Reserch Fellowship under Grant No. 1937968 (KY) and the National Institutes of Health Ruth L. Kirschstein Predoctoral Individual NRSA F31CA200373 (CH). The authors would like to thank Dr. Steven Schomisch for providing pig livers. The authors have no competing interests to declare.

## Data Availability

The raw and processed data required to reproduce these findings are available to download from: http://dx.doi.org/10.17605/OSF.IO/B8DNW.

