## Supplemental Methods and Results for "Conventional polymers may unintentionally refill in vivo with unassociated drugs"

### Supplementary Methods

#### Polymerized Cyclodextrin Microparticle Drug Loading

An alternative drug loading strategy to incorporating drug directly into ISFI polymer solution is to pre-load pCD microparticles with DOX. A high concentration solution of DOX in MilliQ water was made at 5 mg/mL. pCD microparticles were added to loading solution at a ratio of 30 mg pCD to 1 mL solution in 1.5 mL microcentrifuge tubes. This ratio ensured even permeation of loading solution throughout the pCD during drug loading on an end over end mixer for 3 days. At the end of drug loading, particles were washed at least 3 times with DI water, or until the supernatant ran clear, to remove excess unbound drug. Particles were then frozen and lyophilized for 3 days, and drug-loaded particles were obtained as a dry powder.

#### Polymer Solution Preparation and Implant Synthesis

Phase inverting in situ forming implants (ISFIs) were synthesized by forming PLGA solutions in water-miscible N-methyl-2-pyrrolidone (NMP), according to previously published protocols.[18] All polymer solutions were formulated with a 60:40 mass ratio of NMP to combined polymer and drug. Implants containing pCD were made with 10 wt.% pCD at a 60:30:10 mass ratio of NMP:PLGA:pCD. This was the maximum amount of pCD incorporation possible into the ISFIs; above this amount, the solution would not form a single solid matrix. After fully mixing, polymer solution was used within 1 day. Implants were formed by pipetting 40 uL of polymer solution into 10 mL of PBS in glass vials.

### Swelling and Erosion Studies

To understand how the incorporation of pCD affects ISFI formation and physical properties, we quantified swelling and erosion over time. To obtain swelling ratio and erosion, implants were synthesized according to the methods described in *Polymer Solution Preparation and Implant Synthesis* without drug loading. Initial implant weights were recorded, and initial polymer weight was calculated based on polymer solution component ratios (40% polymer). PBS bath solution was changed on a daily basis, and at specified timepoints implants were removed from solution (n=3): time t=2 hours, 4 hours, 1 day, 2 days, 3 days, 5 days, and 7 days. Implants were blotted gently on tissue to remove excess liquid and weighed to obtain wet weight. Implants were frozen in MilliQ DI water at -80ºC and lyophilized overnight. Then mplants were weighed to obtain dry weight. Swelling Ratio was calculated as follows:

$$Swelling\ Ratio = \frac{Wet\ Weight - Dry\ Weight}{Dry\ Weight}$$

Erosion (% Polymer Mass Remaining) was calculated as follows:

$$Erosion = \frac{Dry\ Weight}{Initial\ Polymer\ Weight} * 100\%$$

### Drug Loading Capacity *in vitro*

The maximum amount of drug that could be loaded into “empty” ISFIs was evaluated to get a measure for their potential to be loaded with drug. ISFIs without drug were placed in high concentration drug loading solutions, and the amount of loaded drug was determined by dissolution and solvent extraction. Briefly, implants without drug (PLGA-only, PLGA-pCD) were synthesized and stored on an incubated shaker at 37 °C and 100 rpm for 3 days prior to use. Drug loading solutions were made 0.2, 1, and 5 mg/mL of DOX in PBS. Implants were incubated in loading solutions at 37 °C and 100 rpm for 3 days, removed from the solutions to end loading, rinsed briefly in DI water, and then patted dry with tissue (Fig. 1A). To quantify the amount of loaded DOX, implants were dissolved in DMSO, and overall drug fluorescence was quantified on a multimode microplate reader (Synergy, H1 Microplate Reader). Fluorescence was compared to a standard curve of known DOX concentrations to convert fluorescence to drug mass.

#### Drug Release Studies *in vitro*

In application, ISFIs would be synthesized with an initial load of drug, therefore we sought to characterize how incorporation of pCD as well as how different drug loading strategies would affect drug release profiles. Implants were synthesized with initial loading of 1 wt.% DOX: Polymer solution for PLGA ISFI were made with 60:39:1 NMP:PLGA:DOX mass ratio, and polymer solution for PLGA-pCD ISFI were made with 60:29:10:1 NMP:PLGA:pCD:DOX mass ratio. ISFI release vials were stored on an incubated shaker (37 °C at 100 rpm). During the first 24 hours of release after implant formation, 1 mL of release solution was sampled at each timepoint, and then 1 mL fresh pre-warmed PBS was added back to maintain infinite sink conditions. After 1 day of drug release, 1 mL of release was sampled and then the remaining 9 mL of release solution was replaced with 10 mL fresh pre-warmed PBS. Quantification of DOX in release aliquots was determined by measuring the fluorescence (excitation: 498 nm, emission: 590 nm) and comparing to a standard curve of known masses on a Synergy H1 Microplate Reader (BioTek, Winooski, VT).

#### Measurement of DOX-loaded ISFI Bioactivity

As an initial study of the bioactivity of PLGA-pCD implants, drug release aliquots from the drug release studies above were used in a cell viability (MTS) assay against N1-S1 rat liver hepatoma cancer cells (ATCC, Manassas, VA). Bioactivity over time depicted as % growth inhibition of N1-S1 cells treated with release aliquots compared to cells grown in 100% growth medium (IMDM with 10% fetal bovine serum). DOX standards in complete growth medium were made at concentrations 0.005-100 ug/mL. N1-S1 cells were seeded at 5000 cells/well in 96 well plates, and diluted 1:1 with DOX standards and release aliquots to a total well volume of 100 uL. Standards and release aliquots were added in triplicate; all wells on the edge of the plate were filled with PBS to reduce evaporation. Cells were incubated for 48 hours, then 20 uL of MTS solution were added to each cell-containing well, covered with foil and incubated for 1-4 hours. At 30 minutes, 1, 2, and 4 hours, plate absorbance was measured at 490 nm on a Synergy H1 Microplate Reader (BioTek, Winooski, VT). Cell Proliferation was normalized by dividing by the negative control, proliferation measured for cells in complete growth medium. Growth Inhibition was calculated as follows:

$$Growth\ Inhibition\ (\%) = \frac{Negative\ Control - Proliferation}{Negative\ Control} * 100\%$$

### Statistical Testing

Student's t-test was used for statistical testing. All data is shown as mean ± standard deviation.

### Supplementary Figures and Tables

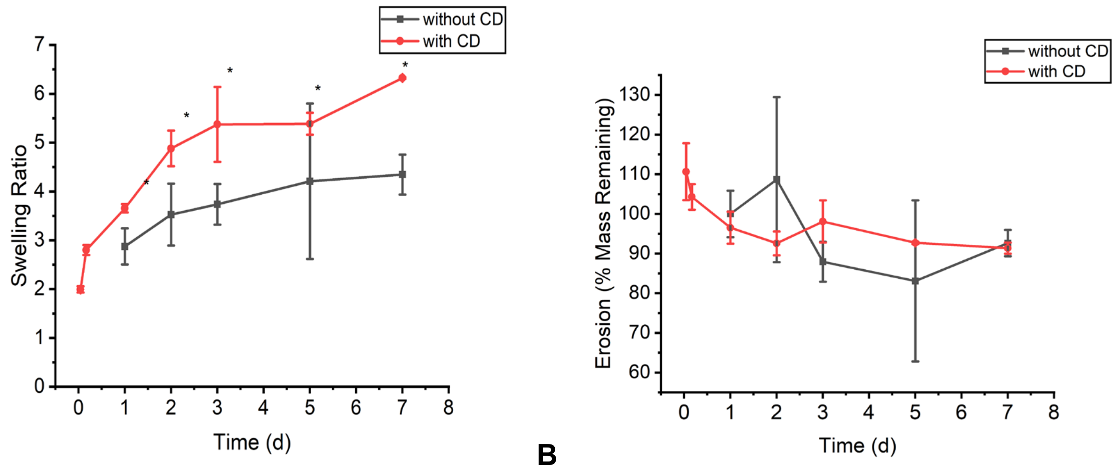


**A** **B**

**Figure S1**. **Differing characteristics of PLGA vs PLGA-pCD ISFI shown through swelling and erosion studies.** For days 1-7, PLGA-pCD ISFI exhibited significantly greater swelling compared to PLGA ISFI, however, there was no significant difference in erosion. * indicates $p<0.05$.

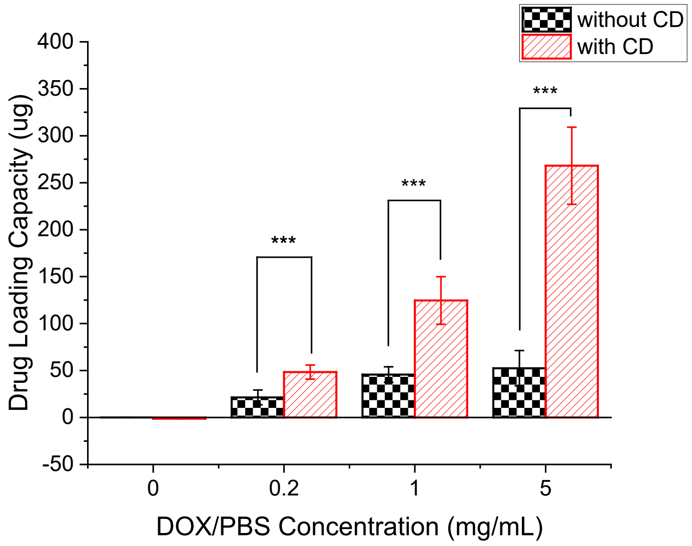


**Figure S2**. **Drug loading of DOX into ISFI at varying loading concentrations, without normalization by initial implant weights.** This shows that for ISFIs synthesized with *similar initial volume*, PLGA-pCD ISFI is capable of loading with significantly more DOX compared to PLGA ISFI – a more than 5-fold increase. pCD microparticles are denser and heavier than the PLGA that was used, therefore PLGA-pCD are heavier. *** indicates $p<0.001$.

**Table S1. Fold increase in DOX loading calculated as the ratio of loaded DOX in PLGA-pCD to PLGA-only, without normalization by initial implant weights.** At lower DOX refilling concentrations, PLGA-pCD implants refilled with over 2 times the amount of DOX. At the highest refilling concentration, PLGA-pCD implants notably refilled with 5 times the amount of DOX, compared to PLGA-only implants.

| **Loading Solution (mg/mL)** | **Fold Difference (PLGA-pCD:PLGA)** |
|---|---|
| 0.2 | 2.264 |
| 1 | 2.726 |
| 5 | 5.110 |

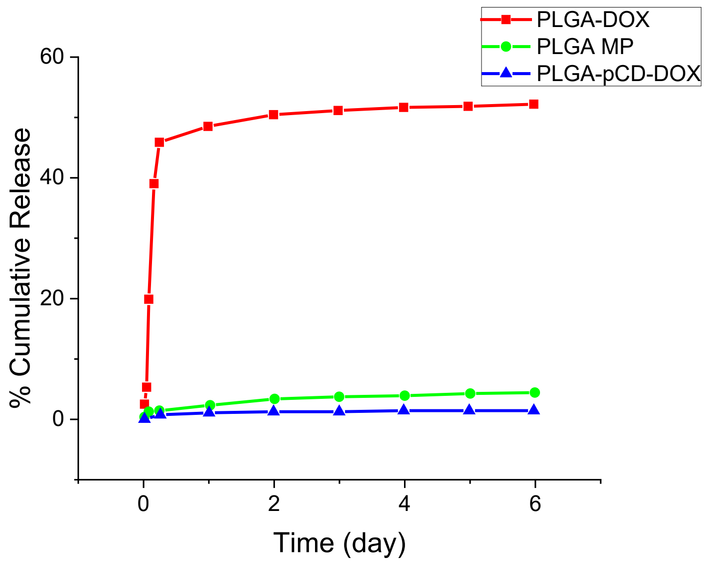


**Figure S3. Drug release profiles obtained from different drug loading strategies:** 1) PLGA ISFI loaded with 1 wt.% DOX, 2) pCD microparticles loaded with DOX (PLGA MP), and 3) PLGA-pCD ISFI with 10 wt.% pCD microparticles loaded with DOX (PLGA-pCD-DOX). pCD microparticles were synthesized following the methods described in Polymerized Cyclodextrin Microparticle Synthesis. pCD microparticles were loaded with DOX by incubating in 5 mg/mL DOX/water solution at 30 mg microparticles per 1 mL of solution on an end over end mixer for 3 days. DOX-loaded particles were washed several times with DI water to removed excess unbound drug, then frozen in MilliQ water and lyophilized for 3 days to obtain dry DOX-loaded particles in powder form. Drug release from PLGA-DOX resulted in a high initial burst release, followed by a slower lag phase, as expected. Drug release from PLGA MP was much more controlled, with approximately 5% cumulative drug released at day 6. Drug release from PLGA-pCD-DOX was even more controlled and slower than PLGA MP for the period of 6 days, likely due to DOX needing to diffuse through the PLGA matrix in addition to overcoming affinity with pCD in order to be released.

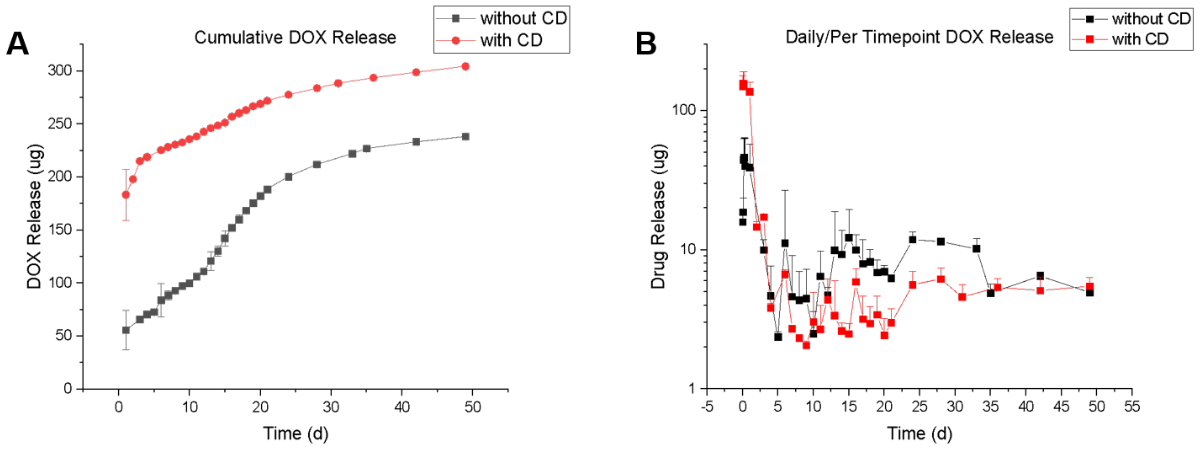


**Figure S4**. **Drug Release Studies from ISFI initially loaded with a standard amount of DOX (1 wt.% loading)**. **A**) Calculated cumulative release shows the overall amount of drug released. PLGA-pCD ISFI released more drug than PLGA ISFI over the course of 50 days. **B**) Drug release replotted as Daily/per Timepoint release on a semilog scale to show release rates over time. Incorporation of free DOX within PLGA-pCD ISFI actually resulted in much greater burst release compared to free DOX in PLGA ISFI (**Fig. S3**). Incorporation of pCD microparticles increased initial burst (0-24 h) and slowed down subsequent drug release. Release profiles followed typical kinetics from degradable devices: initial high rate of burst release followed by slow-rate lag phase release, finally followed by degradation-based released. [citation?] Addition of CD particles increased initial burst release in the first 24 hours by more than 300%. This was due to rapid implant formation as a result of accelerated solvent exchange with the presence of hygroscopic pCD.

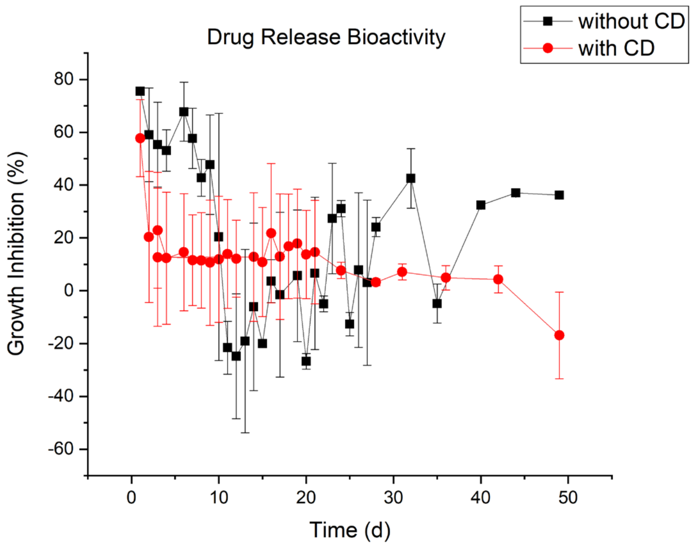


**Figure S5**. **Bioactivity (anti-cancer cell activity) over time of PLGA-only and PLGA-pCD ISFI with 1 wt.% DOX initial loading depicted as % Cell Growth Inhibition**. Implants with CD maintained 10% growth inhibition for 3 weeks. Given that cells were treated with a fraction of each release aliquot due to MTS assay parameters, it is likely that the growth inhibition observed is an underestimation of the full effect of the entire release aliquot. The amount of drug released from single implants were insufficient to inhibit the chosen number of cells beyond 10%. The efficacy of these ISFI can be modulated by increasing the size and number of ISFI.
